# Assessing Comparative Microbiome Performance in Plant Cell Wall Deconstruction Using Multi-‘omics-Informed Network Analysis

**DOI:** 10.1101/2022.01.07.475446

**Authors:** Lauren M. Tom, Martina Aulitto, Yu-Wei Wu, Kai Deng, Yu Gao, Naijia Xiao, Beatrice Garcia Rodriguez, Clifford Louime, Trent R. Northen, Aymerick Eudes, Jenny C. Mortimer, Paul Adams, Henrik Scheller, Blake A. Simmons, Javier A. Ceja-Navarro, Steven W. Singer

## Abstract

Plant cell walls are interwoven structures recalcitrant to degradation. Both native and adapted microbiomes are particularly effective at plant cell wall deconstruction. Studying these deconstructive microbiomes provides an opportunity to assess microbiome performance and relate it to specific microbial populations and enzymes. To establish a system assessing comparative microbiome performance, parallel microbiomes were cultivated on sorghum *(Sorghum bicolor* L. Moench) from compost inocula. Biomass loss and biochemical assays indicated that these microbiomes diverged in their ability to deconstruct biomass. Network reconstructions from time-dependent gene expression identified key deconstructive groups within the adapted sorghum-degrading communities, including *Actinotalea, Filomicrobium,* and Gemmanimonadetes populations. Functional analysis of gene expression demonstrated that the microbiomes proceeded through successional stages that are linked to enzymes that deconstruct plant cell wall polymers. This combination of network and functional analysis highlighted the importance of cellulose-active Actinobacteria in differentiating the performance of these microbiomes.

## Introduction

Plant cell walls are complex structures that primarily contain the polysaccharide polymers cellulose, hemicellulose, and pectin as well as the aromatic polymer lignin^1^. The primary cell wall of grasses, such as sorghum *(Sorghum bicolor* L. Moench), is a thin layer consisting of cellulose and the hemicellulose xylan, and a small amount of pectin. The thicker secondary cell wall, deposited after plant cell growth ceases, contains cellulose, lignin, and hemicellulose. Chemical and biological deconstruction of plant cell walls to release the sugars and aromatics in the biomass is of great current interest for their subsequent conversion to biofuels and bio-based chemicals^2–4^. For biological deconstruction, microorganisms, including filamentous fungi, bacteria, and protists employ an armamentarium of enzymes that systematically deconstruct the plant cell wall^5–7^. These include hydrolytic and oxidative enzymes that deconstruct the polysaccharides and radical-based oxidative enzymes that deconstruct lignin^8–10^.

Though most understanding of biological cell wall deconstruction has been obtained from isolates, microbiomes that break down cell walls have emerged as new sources of microbes and enzymes^11–15^. These microbiomes feature successional structures that are linked to the mechanism of depolymerization in the cell wall^16^. Microbiomes that digest plant cell walls are readily cultivated from inocula rich in deconstructive microbes, like compost and rumen^17,18^. These cultivations have yielded microbiomes with reproducible structures and community dynamics, linking plant polymer deconstruction to individual microbes and enzymes. Development of parallel consortia from heterogeneous inocula leads to variations in microbiome structure, often referred to as founder effects, that may influence microbiome performance^19^. Therefore, the cultivation of parallel consortia is a promising strategy to link the structure and dynamics of biomass-deconstructing microbiomes. These comparisons may identify key contributors to the deconstruction of cell wall components that differentiate the microbiomes.

Here, parallel microbiomes with different community structures were cultivated with sorghum biomass as the sole carbon source. The performance of these distinct microbiomes was compared in growth on forage sorghum varieties and this performance linked to specific populations by network and functional analyses of time-resolved metatranscriptomics.

## Materials and Methods

### Sample Collection and Biomass Preparation

Green waste compost was collected from the City of Berkeley (https://www.cityofberkeley.info/freecompost/) and transported to the lab at room temperature. Compost was sieved and stored at 4°C prior to use. Untreated ground *Sorghum bicolor* L. (25 mm particle size) was obtained from Idaho National Laboratory and washed, autoclaved, and dried in a 50C oven. The wild type and *brown midrib-6×12 (bmr-6×12)* forage sorghum were grown and harvested as previously described^20^. The forage sorghum samples were also washed, autoclaved, and dried as described above. Moisture content was measured using a moisture analyzer (Mettler Toledo Moisture Balance HB43-S).

### Enrichment/Priming (Tier 1)

Green waste compost (0.1 g), 50 mL of M9TE^21^ (pH 6.5), and 0.5 g of sorghum were inoculated in 250 mL baffled Erlenmeyer flasks. Three parallel incubations, along with a negative control without inoculant, were incubated at 50 °C at 200 rpm and adjusted for evaporation using filter-sterilized deionized water every 2-3 days. Passages were conducted every 2 weeks (Day 14, 28, 42, and 56) by transferring 2 mL of culture to a new set of flasks. At the end of each passage, pH was measured and 500-μl aliquots were collected and centrifuged to separate pellet and supernatant fractions. DNA was extracted from the pellet fraction and sent for 16S rRNA gene and metagenomic sequencing. Additionally, for the final passage (Day 56), 3,5-dinitrosalicylic acid (DNS) assays^22^ and nanostructure-initiator mass spectrometry (NIMS)^23^ assays were performed on the supernatant fraction and the remaining material was filtered using Miracloth (Millipore Sigma, Burlington, MA, USA) and dried at 50 °C to determine the biomass dry weight.

### Dynamics (Tier 2)

At Day 56, each of the three communities from Tier 1 *(comm1, comm2,* comm3) were used to inoculate a second series of flasks (Tier 2). Two milliliters of each sorghum-deconstructing microbiome (SDM) from Passage 4 (Day 56) was used to inoculate triplicate flasks containing 50 ml M9TE (pH 6.5), and 0.5 g of either the parent forage sorghum or *bmr6* x *bmr12* stacked mutant^24^. Triplicate flasks along with a control were incubated at 50 °C, 200 rpm for 2 weeks. At each timepoint (Day 2, 5, 7, 9, 12, and 14), flasks were adjusted for evaporation, measured for pH, and sampled for nucleic acid extraction. Five hundred-microliter samples were centrifuged for 5 min at 14,000 x g and pellets used for DNA/RNA co-extraction. After 14 days, 500 μl of media were centrifuged, and supernatant used for DNS assays and NIMS analysis. NIMS analysis was performed as described in detail elsewhere^25^. Briefly, a 2 μL aliquot of supernatant was transferred into a vial containing 6 μL of 100 mM glycine acetate, pH 1.2, 0.5 μL of a 5.0 mM aqueous solution of [*U*]-^13^C glucose, 2 μL of CH3CN, 1 μL of MeOH, 1 μL of solution probe (100 mM in 1:1 (v/v) H_2_O:MeOH), and 0.1 μL of aniline. The mixture was incubated at room temperature for 16 hours. NIMS analysis was performed using a Bruker UltrafleXtreme MALDI TOF/TOF mass spectrometer. In each case, 0.2 μL of the quenched reaction sample was spotted onto the NIMS surface and removed after 30 seconds. Signal intensities were identified for the ions of the tagging products and ~4000 laser shots were collected. Residual biomass was filtered through Miracloth and a subsample of 100 mg used for lignin quantification using the Acetyl Bromide Soluble Lignin (ABSL)^26^ assay and the rest dried to determine dry weight. DNA from Day 14 was used for metagenome sequencing, while RNA from each sampling point submitted for metatranscriptome sequencing as described below.

### DNA/RNA Extraction for Metagenomics and Metatranscriptomics

DNA and RNA were co-extracted from 500uL of SDM pellets as previously described^27^ using a modified CTAB extraction buffer consisting of equal volumes of 0.5 M phosphate buffer (pH 8) in 1 M NaCl and 10% hexadecyltrimethylammonium bromide (CTAB) in 1 M NaCl. Briefly, tubes containing 500 μl of SDM pellet, 0.5 mL of modified CTAB extraction buffer, 50 μl of 0.1 M ammonium aluminum sulfate and 0.5 mL of phenol:chloroform:isoamyl alcohol (25:24:1) were bead-beaten at 5.5 m/s for 45 s in a FastPrep instrument (MP Biomedicals, Solon, OH, United States). Following bead-beating, tubes were centrifuged at 16,000 × g for 5 min at 4 °C. The supernatant was transferred to a new tube containing an equal volume of chloroform:isoamyl alcohol (24:1), vortexed, and centrifuged again. The supernatant was transferred into a new tube containing 1 ml of polyethylene glycol 6000 solution and 1 μl of linear acrylamide and incubated at room temperature for 2 h. Each sample was extracted a second time by adding 0.5 ml of modified CTAB extraction buffer to the original Lysing Matrix E tubes and repeating the steps from beadbeating onwards. The first and second extractions were centrifuged at 16,000 × g for 10 min at 4 °C. The pellets (two per sample) were washed with 0.5 ml of cold 70% ethanol, dried, and combined in 50 μl of RNase-Free water. Purification was carried out using the AllPrep DNA/RNA Mini Kit (Qiagen, Valencia, CA, United States) according to manufacturer’s instructions. DNA and RNA were eluted in 60 μl and 30 μl of RNase-Free water, respectively. Concentrations were measured by Qubit fluorimeter (Invitrogen, Carlsbad, CA, United States) and quality was assessed by BioAnalyzer (Agilent).

### Characterization of bacterial communities with amplicon sequencing

Triplicated amplicon libraries were prepared using 3 ng of DNA per reaction and the primers 515F and 806R modified with Illumina sequencing adapters and barcodes. Libraries were pooled in equimolar concentrations and sequenced on the MiSeq platform using the Miseq Reagent kit v3. Sequences were demultiplexed based on their unique barcodes and trimmed to the same length. Sequences were dereplicated and sorted by decreasing abundance using USEARCH v11^28^. The dereplicated sequences were denoised, *de-novo* chimera filtered, and zero-radius OTUs (ZOTU) generated using unoise3 from USEARCH v11. Resulting ZOTUs, which are a form of amplicon sequence variants (ASVs), were taxonomically characterized against the Greengenes database gg_16s_13.5 using Sintax (USEARCH v11) with a cutoff of 0.8, and genus as the maximum taxonomic level. Total sequences were mapped against the ZOTUs at a 97% identity and an abundance table was generated that was subsequently transformed into a biom table. ZOTUs were aligned using Clustalw, and the alignment was used to generate a phylogenetic tree with IQ-TREE 2^29^ using the model TIM3+F+I+G4 (identified using model finder) and ultrafast bootstrap approximation (UFBoot) with 1000 replicates. The abundance table, mapping file, and phylogenetic tree were imported to the R software using the Phyloseq package^30^ (version 1.12.2). For community composition analyses (beta-diversity), data was VST-normalized using the DESeq2 package^31^ (version 1.34.0) using a mean fit that was used to calculate a weighted Unifrac distance matrix. The obtained distance matrix was ordinated using multidimensional scaling in Phyloseq. The samples were categorized based on passage and its effect on data variation tested with Adonis (nonparametric permutation multivariate analysis of variance), performed with 1,000 permutations.

### Metagenomic Sequencing and Analysis

Twenty-one DNA samples, 3 from Tier 1 Day 56, and 18 from Tier 2 Day 14, were submitted to the Joint Genome Institute (JGI) for sequencing using Illumina Novaseq platform (150 bp x 2). Individual reads were filtered using JGI’s standard metagenomic analysis pipeline (version 3.4.7 from BBtools version 38.24), corrected using bbcms (version 38.34), and co-assembled using metaSPAdes^32^ (version 3.13.0). Open Reading Frames were predicted from the assembled contigs using MetaGeneMark^33^. Protein domain annotations were predicted using the pfamA-30 and dbCAN-V8 Hidden Markov Model protein domain databases using an e-value of 1 x 10^-5^. Protein categories of interest were screened against the National Center for Biotechnology Information database using BLASTp and dbCAN2’s CAZy database for DIAMOND^34^ (version 0.9.21.122) with an e-value 1 x 10^-5^. The metagenome co-assembly was binned using MaxBin (version 2.2.5) with default parameters, yielding 103 Metagenome Assembled Genomes (MAGs). The most likely taxonomy was predicted using a custom script (getTaxon.pl), which searched the predicted proteins of the individual bins against the NCBI non-redundant (NR) database using DIAMOND (version 0.9.21.122) and processed the hits using the least common ancestor (LCA) algorithm proposed by MEGAN Community edition (version 6.11.0)^35^. Completeness and contamination rates for all MAGs were assessed using CheckM (version 1.0.12). MAGs (and associated genes) with at least 30% completeness and less than 10% contamination were retained for the rest of the analyses. Coverage information for the scaffolds of each MAG was extracted from the calculated coverage data TPM normalized data for each scaffold in the metagenome, and MAG abundances in each replicated sample were calculated as the average TPM coverage value over all the scaffolds in a MAG. The compositional variation of each enriched community was analyzed by quantifying their Local Contribution to Beta Diversity (LCBD) using the R package *adespatial* with the Hellinger dissimilarity coefficient and *p*-value correction using the Holm method. A phylogenetic tree for the MAGs was reconstructed in KBase^36^ based on universal genes defined by Cluster of Orthologous Groups using maximum likelihood. Average Nucleotide Identity between taxonomically related MAGs (genus level) was quantified also in KBase. Annotations for each of the MAGs are provided in Supplementary Data.

### Metatranscriptomic Sequencing and Analysis

Fifty-four RNA samples, from each of the treatments and time points of Tier 2 experiment were also submitted to JGI for metatranscriptomic sequencing using the Illumina Novaseq platform (150bp x 2). Sequenced samples represented triplicated RNA samples from adapted communities incubated with stacked mutant and WT sorghum. The filtered reads were assessed using FastQC (version 0.11.8) and mapped to the metagenome co-assembly using Bowtie2 (version 2.3.4.3). Gene counts were generated using Feature Counts (version 1.6.3) and normalized for both gene length and library size by transcripts per million (TPM), using a custom R script. For metatranscriptome ordination analyses a Bray-Curtis dissimilarity matrix was calculated using R’s Vegan on the raw feature counts table that was first filtered to retain only those genes appearing in at least 5 samples (out of the total 54 samples) and mean count of 10. The resulting table was VST-normalized with DESeq2. The samples were categorized based on time (day), type of biomass (WT and SM), and categorical effects on data variation tested with Adonis (nonparametric permutation multivariate analysis of variance), performed with 1,000 permutations. Average transcriptome abundances per selected MAG were calculated on the TPM-normalized data and are available in Supplementary Data. For differential expression analyses, the feature count data was filtered using the parameters used for the transcriptome ordination analysis, retaining genes appearing in at least 5 samples with a mean count of 10. Differential expression analyses were carried using DESeq2 using a parametric fit. The results filtered for a corrected *p*-value < 0.01 and an absolute log2fold change > 1. Heatmaps showing normalized expression levels per relevant genes were calculated on the DESeq-2 VST-normalized data using R’s *pheatmap* package, and rows arranged based on a Bray-Curtis dissimilarity matrix.

### Network reconstruction

A network was constructed for the transcriptome data based on centred logratio transformed feature counts data^37^. Prior to normalization, the data was subsetted to include genes detected in at least 50% of the total number of samples. Network reconstruction was conducted with the Molecular Ecological Network Analyses pipeline (MENAP, http://ieg4.rccc.ou.edu/mena/) with the following settings: for missing data fill blanks with 0.01 if data have paired values; do not take logarithm as the data was already CLR normalized; use Spearman Correlation similarity matrix; calculate by decreasing cutoff from the top. Random Matrix Theory (RMT) was used to automatically identify the appropriate similarity threshold for network reconstruction^38,39^. The network was visualized in Cytoscape^40^ (version 3.9.0) using Force-Directed graph drawing and colored based on the taxonomic identity of the included MAGs.

## Results

### Microbial community adaptation to grow on sorghum

Green waste compost was used to inoculate three parallel microbiomes which were adapted to grow on sorghum biomass as the sole carbon source for 56 days. Measurement of residual sorghum biomass by Day 56 showed that *comm1* and *comm2* had a 40% reduction in biomass content and *comm3* had a 57% biomass reduction (Supplementary Fig. 1). Amplicon sequencing demonstrated that these microbiomes differentiated into individual communities *(comm1, comm2* and *comm3)* Analysis of community composition showed that the individual microbiomes did not group over time (PERMANOVA: *df*= 3, *F* = 1.59, *p* = 0.21) but rather varied by community (PERMANOVA: *df* = 2, *F* = 4.93, *p* = 0.003, r^2^ = 52.3%) with each following a different trajectory (Fig. 1A). The microbiomes *comm1* and *comm3* were more closely related to each other than *comm2*, which was separated at a considerable distance from the other microbiomes in the ordination plot. The trajectories of these microbiomes suggest that they possess distinct metabolic capabilities and that by Day 56 the community composition had stabilized.

**Fig 1.**
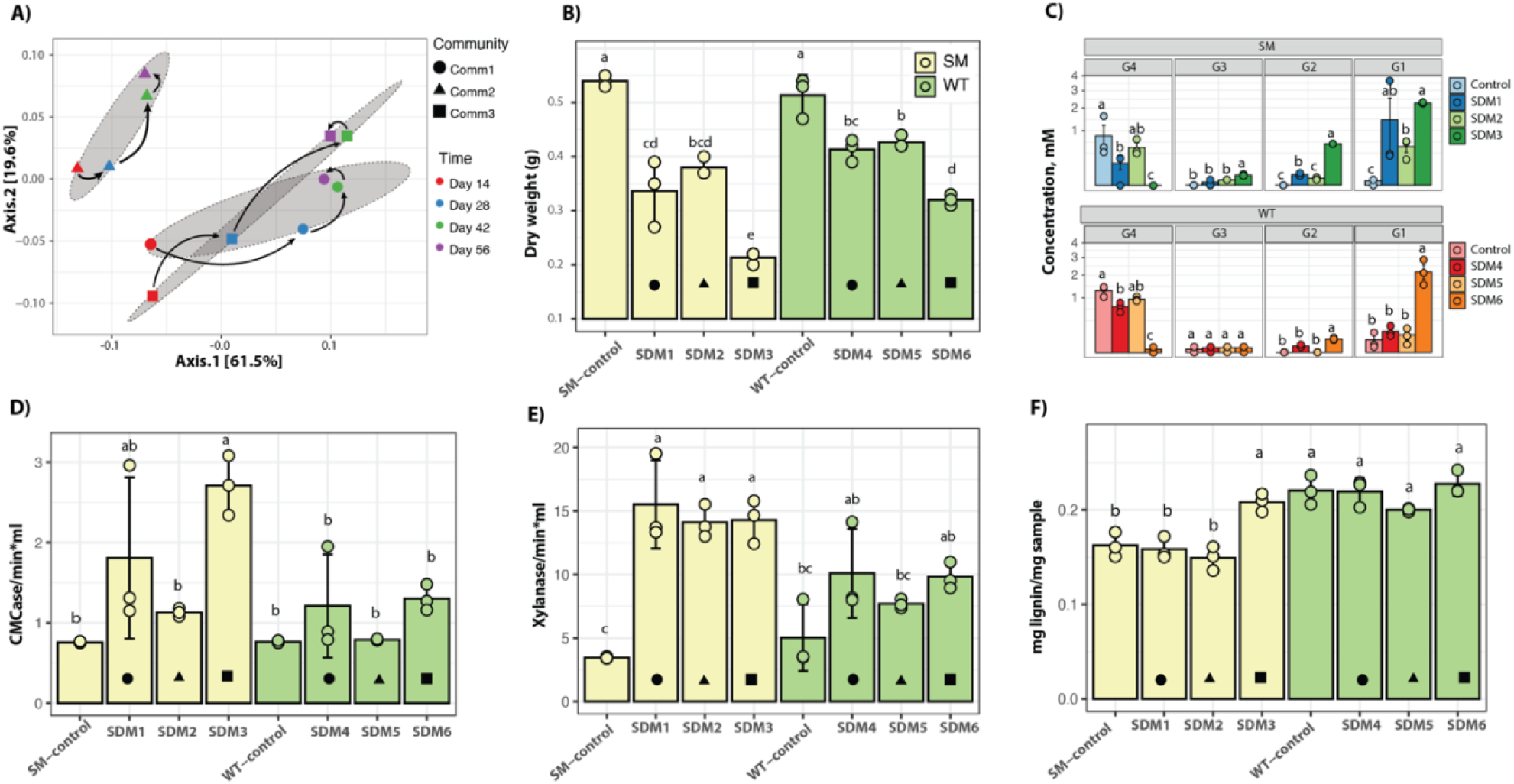
A) Ordination plot for bacterial communities growing on sorghum and analyzed using amplicon sequencing. B) Dry weight; C) NIMS results. Both correspond to end-point analyses after a 14-day incubation. D-E) DNS analysis for CMCase and xylanase activity of adapted communities inoculated to SM and WT sorghum. F) Lignin content from small-scale biomass analysis. The icons within the barplots indicate the Tier1 community used for inoculation of the Tier 2 experiment. Circle – comm1, triangle – comm2, square – comm3. Error bars indicate standard deviation (n = 3). Bars labeled with the same letter are not significantly different (ANOVA and Tukey test; *p* > 0.05).

### Comparative deconstruction of sorghum

The emergence of three distinct microbiomes from the initial green-waste compost inoculum provided an opportunity to compare the performance of parallel microbiomes with different community compositions. We compared the deconstructive abilities of these communities on sorghum varieties with different lignin content and monomeric compositions, to examine the effect of lignin on microbiome performance. Multiple mutants from the lignin biosynthetic pathway have been developed in sorghum, and the *bmr-6*x*12* double mutant was chosen for the parallel experiments^24^. This stacked mutant, in which mutations that affect both the lignin biosynthetic genes cinnamyl alcohol dehydrogenase (*bmr-*6) and caffeic acid O-methyltransferase (*bmr-12*), has lower lignin content and is more easily deconstructed compared to the native sorghum line^41^. Therefore, we compared the corresponding non-mutant sorghum hybrid, referred to as wild type (WT), and the *bmr-6×12* line, referred to as stacked mutant (SM). Microbiomes cultivated for 56 days on forage sorghum were individually inoculated into triplicate cultures containing SM sorghum (SDM1-3) or the WT sorghum (SDM4-6) and cultivated for 14 days. Endpoint measurements of residual dry-weight biomass demonstrated that the communities cultivated on the SM sorghum exhibited the greatest biomass loss. Among the SM communities, SDM3 had a significantly higher average biomass reduction (75% vs control), while SDM6, inoculated using WT-sorghum, exhibited 54% biomass loss compared to the control. SDM3 (SM-treatment) and SDM6 (WT-treatment) are derived from the same *comm3,* and the levels of biomass consumption showed that SDM3 was significantly higher than SDM6 (ANOVA and Tukey test, *p* < 0.01). An analysis of cellotetraose hydrolysis showed that regardless of the type of biomass used as substrate (WT or SM), treatments inoculated with *comm3-derived* microbiome released the highest levels of glucose with SDM3 (SM sorghum) compared to SDM6 (WT sorghum) (SDM3: 2.25 mM, *σ* = 0.04; SDM6: 2.15 mM, *σ* = 0.6) (Fig. 1C). Cellulase and xylanase activity were further investigated using DNS assays and showed the highest enzymatic activity in the *comm1* and *comm3-derived* treatments (SDM1/SDM4 and SDM3/SDM6, respectively). The results also indicated a higher cellulase and xylanase activity in the SM-sorghum treatments compared to WT-sorghum treatment (Fig. 1 D-E).

The changes in biomass composition were further analyzed by measuring relative lignin content compared to uninoculated controls. The residual biomass from the SM communities had a significantly lower lignin content than its WT-counterpart, consistent with the lower levels of lignin in the SM plants versus the WT plants (Fig. 1F). Calculations showed that although not statistically significant (ANOVA and Tukey test, *p* > 0.05) all inoculated treatments had a lower lignin content than their controls, with the exception of the SDM3 treatment, which had an increased amount of lignin in the residual biomass that was statistically significant (Fig. 1F).

### Metagenomic analyses reveal metabolic potential for biomass transformation

A total of 103 metagenome assembled genomes (MAGs) were reconstructed but only 66 that had a completeness above 30% and contamination lower than 10% were considered for downstream analysis. These selected MAGs and their phylogenetic relationships based on universal genes are shown in Supplementary Figure 2. Figure 2A shows the shared and unique reconstructed MAGs in each of the Tier 2 samples and their community sources *(comm1, comm2,* and *comm3)* Inspection of the clustering patterns showed that the composition of Tier 2 samples clustered according to their community sources, as also observed in amplicon-based analysis (Fig 1A). The MAGs separated into five clusters (C1 – C5, Fig 2A). Cluster 1 (C1) represented the communities mostly unique to *comm2* derived samples (SDM2/SDM5) and included *Actinopolymorpha* bin102, *Bacillus* bin91, *Brevibacillus* bin76, 82, and 62 (Average Nucleotide Identity (ANI) = 76%), *Conexibacter* bin85 and 94 (ANI = 78%), *Geobacillus* bin98, *Illumatobacter* bin100, *Microbacterium* bin103, *Mycobacterium* bin99, *Paenibacillus* bin81, *Streptosporangium* bin58, *Thermobacillus* bin92 and 96 (ANI = 77%), and *Ureibacillus* bin93. Cluster 2 (C2) contained bacterial populations shared between *comm2* and *comm3*-derived samples (SDM2/SDM5 and SDM3/SDM6). Cluster 2 included *Actinopolymorpha* bin90, *Bacillus* bin63, *Brevibacillus* bin97, *Paenibacillus* bin101, *Salinispora* bin39 and 64 (ANI = 77%), *Solirubrobacterales* bin89, and *Thermocrispum* bin46. Cluster 3 (C3) represented the populations exclusively shared between *comm3* and *comm1*-derived samples (SDM6/SDM3, and SDM1/SDM4). Cluster 3 populations included *Conexibacter* bin16 and 24 (ANI = 79%), *Inquilinus* bin14, *Mycobacterium* bin18, *Pseudoncardia* bin23, *Salinispora* bin30 and 37 (ANI = 77%). Cluster 4 (C4) represented the core populations among all samples and included *Actinopolymorpha* bin55, *Actinotalea* bin1 and 5 (ANI = 86%), *Aneuribacillus* bin28, *Bacillus* bin60, *Caldibacillus* bin56, *Conhella* bin15, *Dongia* bin26, *Filomicrobium* bin12 and 24 (ANI < 70%), Gemmanimonadetes bin10, *Geobacillus* bin47, *Ornithimicrobium* bin31, *Paenibacillus* bin34, 35, 45, 67, and 69 (ANI = 76% – 78%), *Thermobacillus* bin17, 41, 43, 48, 51, and 53 (ANI = 77% – 89%), *Thermocrispum* bin11, and *Tuberibacillus* bin22. Finally, cluster 5 (C5) included some populations such as the *Rhodospirillales* bin9 and *Salinispora* bin32 which were unique to SDM1/SDM4, and *Thermobacillus* bin96 that was unique to SDM1/SDM6. Other populations in this cluster included *Conhella* bin32, *Thermobacillus* bin49, *Filomicrobium* bin36, *Caldakalibacillus* bin70, and *Paenibacillus* bin42 and 35 (ANI < 70%), all of which were shared between SDM2/SDM5 and SDM1/ SDM4.

**Fig 2.**
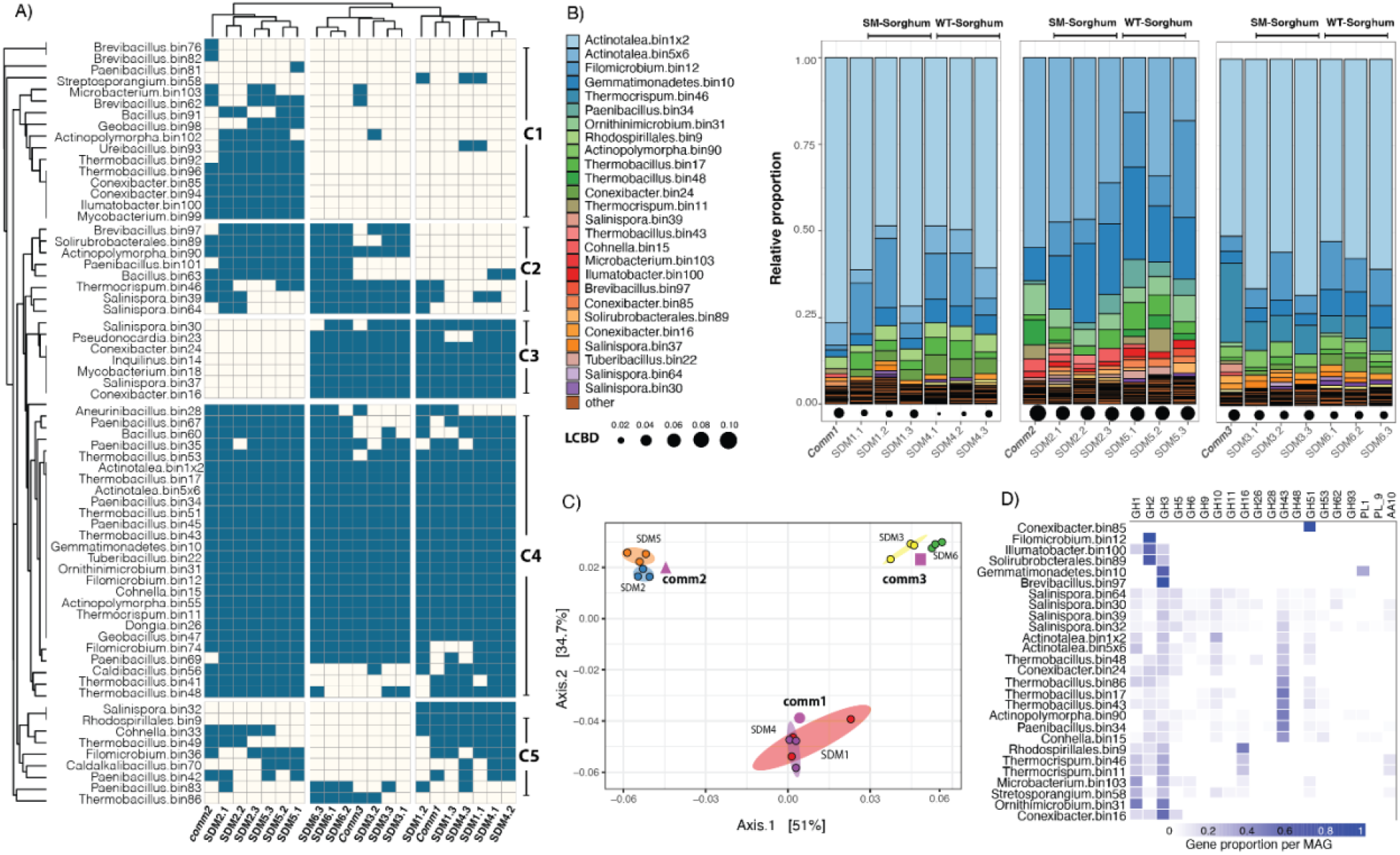
A) Community composition for Tier 2 adapted communities and their source Tier 1 source inoculum. Dendrograms were calculated based on a Jaccard distance matrix. B) Relative proportion of dominant communities calculated from TPM-normalized coverage data. Only populations with a relative proportion above 0.08 are shown in the figure. LCBD = is local contribution to community dispersion calculated with the R package *adespatial;* C) Ordination plot depicting metagenome composition or the Tier 2 adapted communities and their corresponding Tier 1 source inoculum. The ellipses were calculated around barycenters with a confidence level of 0.99 using the *stat_conf_ellipse* function in ggpubr v.0.2.4. D) Gene proportion per MAG for selected GHs.

According to the analysis of coverage distribution of the binned genomes (Fig. 2B), Tier 2 communities were dominated by the *Actinotalea* genome populations (*Actinotalea* bin1 and bin5). *Actinotalea* bin1 contigs accounted for more than 70% of the total contig coverage in SDM1/SDM3 and SMD4/SDM6, while *Actinotalea* bin5 accounted for 41% of the total contig coverage in SDM2 and 24% in SDM5. Highly prevalent MAGs, also identified as part of the cluster 4 (core populations), included populations of *Filomicrobium* bin12, Gemmanimonadetes bin10, *Paenibacillus* bin34*, Ornithiumicrobium* bin31, *Thermobacillus* bin17, 43 and 48*, Thermocrispum* bin46 and 11, and *Cohnella* bin15. Other highly prevalent MAGs were *Brevibacillus* bin97 (C2), *Conexibacter* bin16, 24 (C3), 85 (C1), *Illumiatobacter* bin100 (C1), *Microbacterium* bin103 (C1), *Rhodospirillales* bin9 (C5), *Salinispora* bin37 (C3), 39 (C2), 64 (C2), *Solirubrobacterales* bin89 (C2), and *Thermocrispum* bin46 (C2). Analysis of local contribution to beta diversity (LCBD) showed no significant variation (Holm corrected *p*-values > 0.05) in the composition of the enriched communities when comparing the composition of the Tier2 enrichments and their Tier1 source inoculum (Fig. 2B).

An ordination analysis on the normalized coverage for the contigs of the selected bins (Fig 2C) showed that the different samples clustered together based on their inoculum regardless of biomass type (SM or WT). Furthermore, a permutational analysis of variance showed that the type of inoculum (PERMANOVA: *df=* 2, *F* = 54.9, *p* = 9.9×10^-5^) and type of biomass (PERMANOVA: *df =* 1, *F* = 5.8, *p* = 9.9×10^-5^) had significant effects on metagenomic clustering and explained 84.2% and 4.4% of the observed patterns (Fig. 2C).

Prediction and annotation of genes identified within each MAG showed that the abundant *Actinotalea* bins contained some genes coding for putative glycoside hydrolases relevant for the degradation of polysaccharides. *Actinotalea* bin1 contained GH6 and GH10 genes; while *Actinotalea_*_bin5 had GH5, 6, 10, 43, and 51 genes. On the other hand, other abundant MAGs such as *Actinopolymorpha* bin90, *Conhella* bin15, *Paenibacillus* bin34, *Thermobacillus* bin17 and bin48 contained more of the GHs possibly involved with pectin, hemicellulose, and cellulose degradation (Fig. 2D). Supplementary Figure 3 shows the distribution of relevant GHs among the selected MAGs.

### Sequential degradation of sorghum biomass follows two distinct trajectories

The SDM1 and SDM3 treatments had the same most abundant population (*Actinotalea* bin1) and the highest activities among the Tier 2 microbiomes. Therefore, we performed an indepth comparison of time-dependent gene expression patterns in these microbiomes to identify similarities and differences in expression patterns, focusing on genes for deconstruction of plant polymers. We also performed a comparison between SDM3 and SDM6 to see if the sorghum substrate had any effect on gene expression patterns.

An ordination analysis of the metatranscriptome showed that the three selected enrichments (SDM1, SDM3, and SDM6) followed two distinct trajectories (Fig. 3A). Similar to the metagenome analysis, the metatranscriptomes clustered based on their initial inoculum and shifted gradually over the course of 14 days. SDM3 and SDM6 followed a similar 2-week trajectory, despite having different types of sorghum biomass. SDM1 followed a different trajectory from SDM3 and SDM6, but also exhibited gradual shifts in overall activity, indicative of sequential changes in community structure. A permutational analysis of variance further indicated that type of inoculum (Df = 2, F=40.33, *p* = 9.9×10^-5^) and time (Df = 5, F= 21.59, *p* = 9.9×10^-5^) each had a significant effect on metatranscriptome trajectory, explaining 40.3% and 21.5% of the observed variation, respectively (Fig. 3A). The analysis of variance also indicated that the type of biomass (WT and SM) did not have a significant effect on metatranscriptome trajectory. Based on these results, we chose to focus our analyses on the characterization of SDM1 and SDM3.

**Fig. 3.**
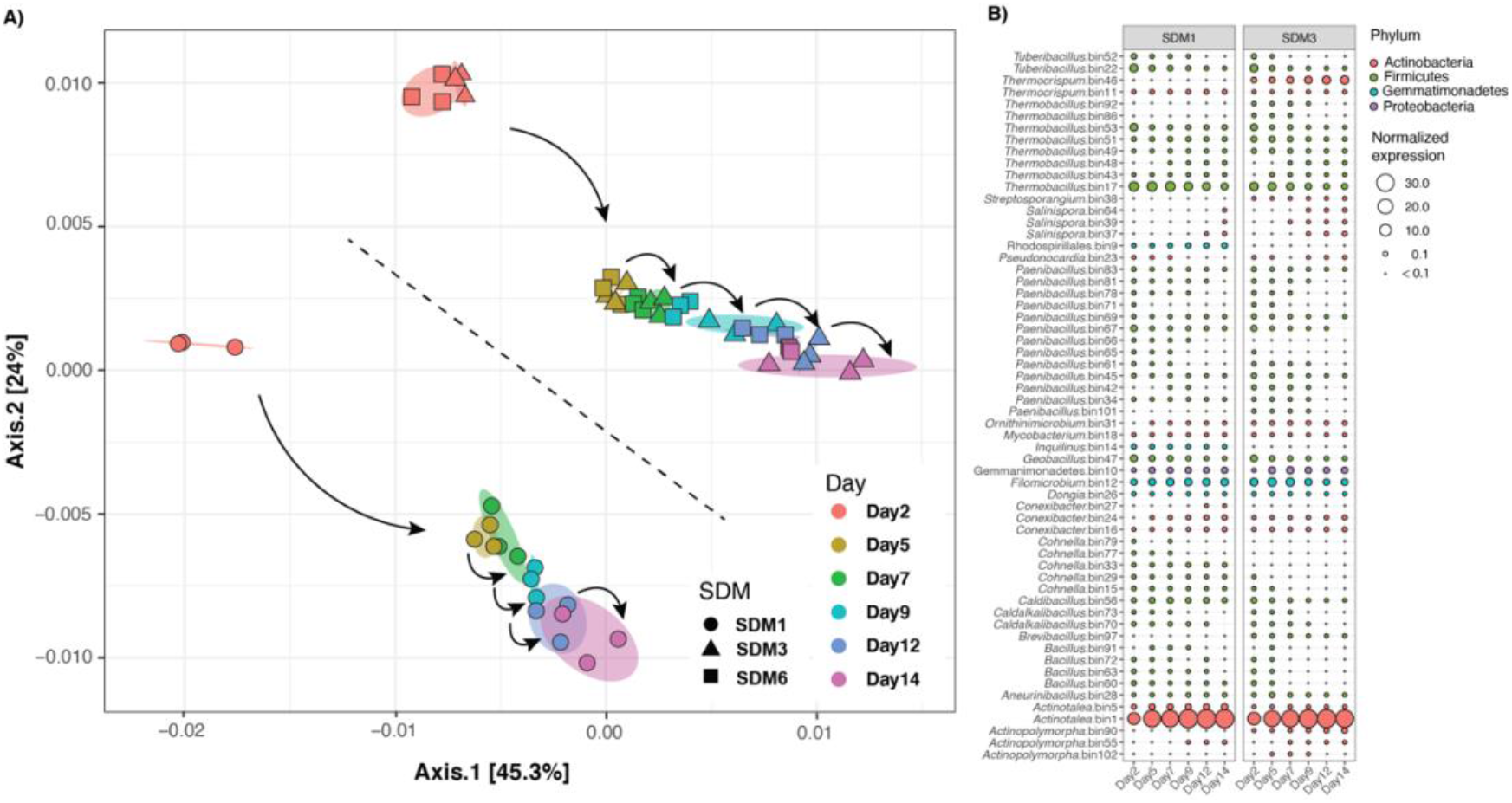
A) Ordination biplot depicting the trajectory of metatranscriptomes for the adapted communities growing on SM and WT sorghum. The ellipses were calculated around barycenters with a confidence level of 0.99 using the *stat_conf_ellipse* function in ggpubr v.0.2.4 B) Average TPM-normalized transcriptome abundance per MAG over the 14-day incubation.

An analysis of the normalized abundance of transcriptomes for each reconstructed MAG indicated that the *Actinotalea-*bin1 was the most active organism in the enrichments across sampling times (Fig. 3B). Other highly active bins included *Thermobacillus* bin17, *Filomicrobium* bin12, *Thermocrispum* bin46, *Gemmatimonadetes* bin10, *Thermobacillus* bin53 and 51, *Tuberibacillus* bin22, *Geobacillus* bin47. Genome bins that were more active in SDM1 included *Actinotalea* bin5, *Rhodospirillales* bin9, and *Inquilinus* bin14, while *Actinopolymorpha* bin90, *Paenibacillus* bin42, and *Thermobacillus* bin49 were more active in SDM3 (Fig. 3B).

Random Matrix Theory (RMT)-based network analysis was performed to define putative interactions among the networked populations and to further explore transcriptome dynamics^38,39^. Figure 4A depicts the reconstructed network based on metatranscriptome expression profiles. Each MAG in the network is colored showing that bacterial populations identified as highly abundant in the metagenome and with high expression levels in the metatranscriptome formed highly connected clusters within the network. The reconstructed network (Fig. 4A) consisted of 22,887 nodes (networked genes) and 5,018,619 links with correlation values between 0.9 – 1.0, and 164 large modules (>10 connected nodes). Cluster isolation by reconstructed MAG with linked neighbors representing co-expression patterns defined potential pairs of interacting bacterial populations. These patterns showed that populations represented by *Actinotalea* bin1, *Actinotalea* bin5, *Filomicrobium* bin12, and *Gemmatimonadetes* bin10 were highly interconnected and likely interacted directly with each other in the metatranscriptomes (Fig. 4B). Because of their conservation in all the microbiomes, high level of abundance and activity (Fig. 2B and 3B), and the direct interconnections between these four MAGs (Fig. 4B) we defined these bins as key populations within the adapted communities. Mapping of differential expression (log2fold change for genes with *p* < 0.01) onto the network showed that *Actinotalea* bin5 was significantly more active in SDM1 during the 14-day incubation (Fig. 4B). We also observed that *Actinotalea* bin1, *Filomicrobium* bin12 and *Gemmatimonadetes* bin10 were more active in SDM3 than in SDM1 from Day 5 to Day 9. The significantly higher activity of these three central bins remained through the 14-day incubation for *Actinotalea* bin1 and declined first for *Gemmatimonadetes* bin10 by Day 12 and then for *Filomicrobium* bin12 by Day 14 (Fig. 4C).

**Fig. 4.**
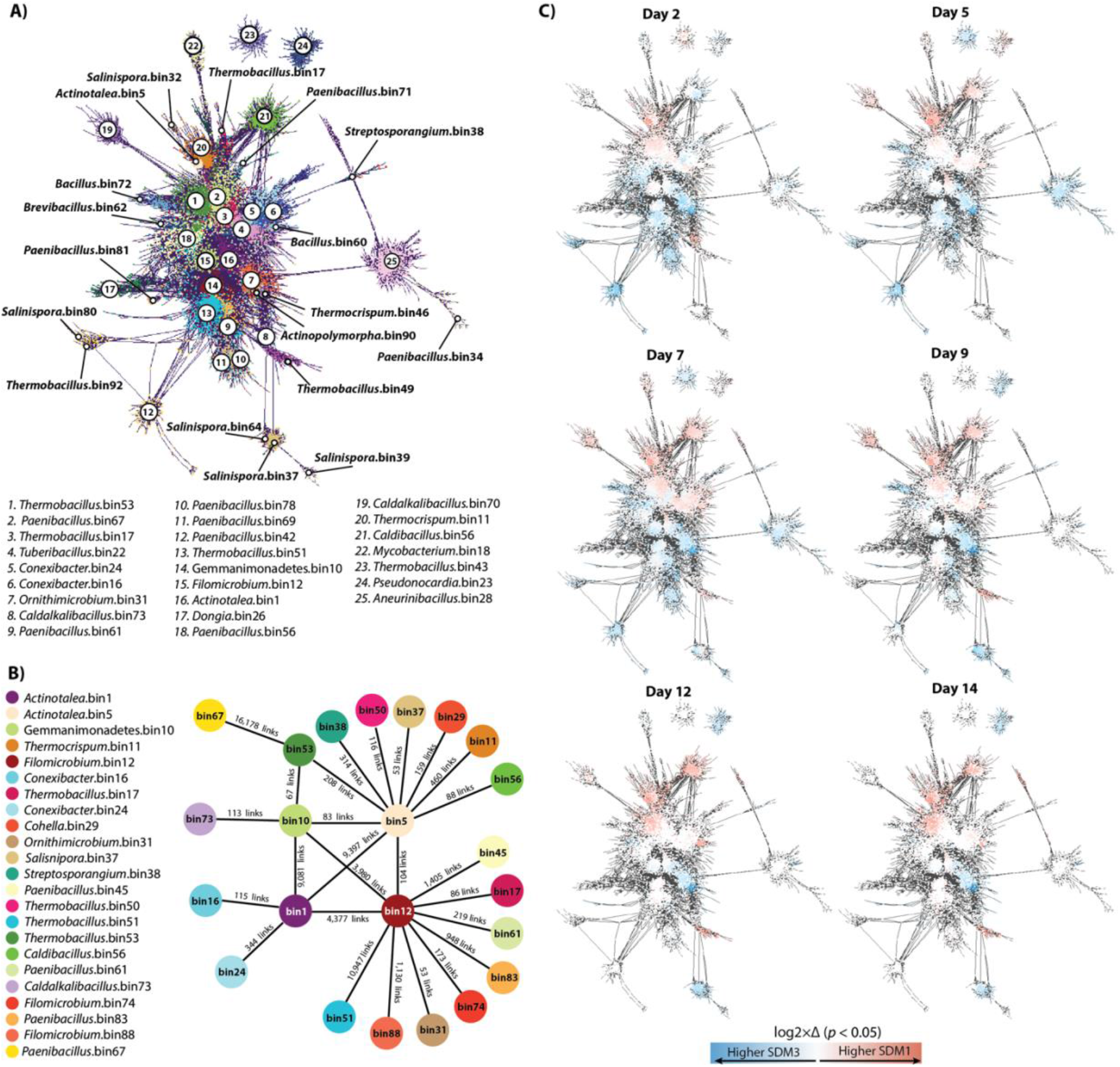
A) RMT-based network reconstructed based on the 14-day metatranscriptome profiles of SDM1 and SDM3 samples. Only significant links with a correlation above 0.9 were retained in the network. B) Illustration of putative population interactions derived from the RMT-network. MAGs connected to the central four MAGs were retained only if connecting by 50 or more links (arbitrary value). C) Differential expression patterns for genes with a log2-fold change higher than 1 and lower than −1 with a *p*-value < 0.01.

One-to-one putative interactions between these four central MAGs with other members of the adapted community were also predicted from the network (Fig. 4A and B). *Actinotalea* bin5 and *Filomicrobium* bin12 had direct connections with a larger number of MAGs than *Actinotalea* bin1 and *Gemmanimodetes* bin10. Populations directly linked with *Actinotalea* bin5 included *Thermocrispum* bin11, *Conhella* bin29, *Salinispora* bin37, *Streptosporangium* bin38, *Thermobacillus* bin50, *Thermobacillus* bin53, *Caldibacillus* bin56. Filomicrobium bin12 on the other hand, had direct links with *Thermobacillus* bin17, *Ornithimicrobium* bin31, *Paenibacillus* bin45, *Thermobacillus* bin51, *Paenibacillus* bin61, *Filomicrobium* bin74, *Paenibacillus* bin83, and *Filomicrobium* bin88. Aside from their connections with the other central MAGs, *Actinotalea* bin1 was found as linked with *Conexibacter* populations bin16 and 24, while *Gemmatimonadetes* bin10 was linked to *Thermobacillus bin*53 (also connected with *Actinotalea* bin5) and with *Caldalkalibacillus* bin73.

We also explored the network associations of MAGs that represented likely key contributors to the process of lignocellulose degradation given their genetic makeup (Supplementary Fig. 1) and high expression levels (Fig. 3B). *Paenibacillus* bin67 was another MAG of interest as it encodes for GHs potentially contributing to the degradation of pectin (GH2, GH43) and hemicellulose (GH10, GH51). *Paenibacillus* bin67 was highly connected with *Thermobacillus* bin53, which contained genes encoding for a wide array of putative GHs including those from families 2, 5, 10, 16, 28, 43, 51, 53, the carbohydrate esterase CE8, and PL1 and PL9 (Supplementary Fig. 2). *Thermobacillus* bin53 was also linked to with *Actinotalea* bin1 and bin5, likely acting as a connection between the dominant *Actinotalea* populations and the rest of the communities.

Another likely prominent group in the process of polysaccharide degradation was the *Salinispora* populations. Three of these MAGs (bin37, 39, and 64) were detected forming a discrete cluster showing high levels of transcriptomic activity in SDM3 from Day 7 to Day 14 (Fig. 4A and 4B). Among these three MAGs, *Salinispora* bin39 and 64 contained a wide arsenal of glycoside hydrolases including GH2, GH5, GH6, GH9, GH10, GH11, GH16, GH43, GH48, GH51, GH53, and PL9 (only bin39); and GH62 and GH93 (only in bin64) (Supplementary Fig. 2).

Detailed exploration of the normalized transcriptome expression profiles indicated that degradation of the primary cell wall was likely initiated by the activity of microorganisms producing enzymes for pectin degradation in a process that was significantly higher in SDM1 than in SDM3 (Wilcox pairwise comparison, *p* < 0.01) and that continued steadily over the 14 days of incubation (Fig 5A). Pectin-degrading expression profiles were separated into four main clusters (Fig. 5D). Cluster 1 (C1) included pectin-degrading genes that were significantly highly expressed in SDM1 and SDM3 *(p* < 0.01, log2fold > 1); C2 and C4 composed by genes highly expressed in SDM3; and C3 genes significantly highly expressed in SDM1. Based on the observed patterns of expression in these clusters, pectin degradation in both treatments was driven by the high levels of expression of GH43 and GH78 from *Actinotalea* bin1 and *Filomicrobium* bin12, respectively. Two main populations of Firmicutes controlled pectin degradation at the start of the incubation in SDM1 including *Thermobacillus* bin53 and *Paenibacillus* bin67 through the expression of most of the genes shown in C1 and C3 at significantly higher levels than in SDM3 *(p* < 0.01, log2fold > 1). Other contributors to the process of pectin degradation in SDM3 were *Thermobacillus* bin51 and Rhodospirillaes bin9, that expressed GH2 and GH43 (in bin51 only) through the whole incubation. Initial drivers of pectin degradation in SDM3 included *Thermobacillus* bin51 and *Filomicrobium* bin12 (C2) and *Salinispora* bin39 (C4) expressing GH78, and *Paenibacillus* bin67 (C2) expressing GH43. *Actinopolymorpha* bin90 (C4) was also among the main contributors to pectin degradation in SDM3 through the expression *(p* < 0.01, log2fold > 1) of GH2, GH43, GH78, and GH93 together with *Thermocrispum* bin46 expressing PL9 and GH2.

**Fig. 5.**
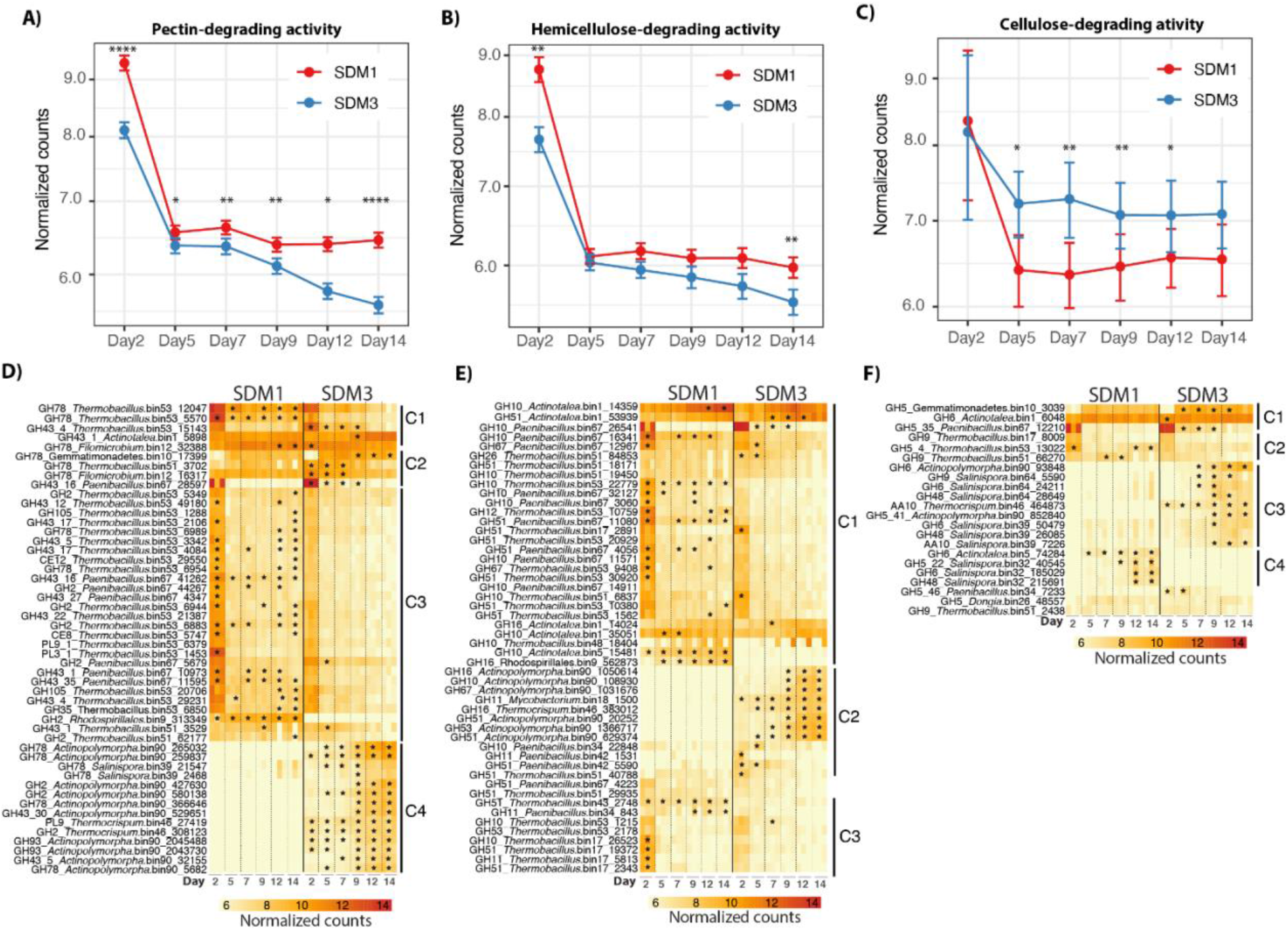
Top panel shows the average trajectories of expression for each of the categories, A) pectin, B) hemicellulose, C) cellulose. Bottom panels depict the different groups of lignocellulose degrading bacterial populations and corresponding gene expression patterns, D) pectin, E) hemicellulose, F) cellulose. Stars indicate the time points at which gene expression was significantly higher than in the opposite treatment (*p* < 0.01, log2fold > 1). GH43 were classified as pectin/degrading enzymes, though this family also cleaves arabinoxylan bonds in hemicellulose^42^.

Hemicellulose deconstruction gene expression dynamics resembled the pectin dynamics as indicated by the higher levels of hemicellulose-deconstructing gene expression in SDM1 than in SDM3 (Fig. 5B). It is likely that *Actinotalea* bin1, the most abundant bacterial population, initiated and maintained the process of hemicellulose deconstruction in both treatments given the high expression levels of the GHs from the families 10, 51 and 16 from this MAG (C1, Fig. 5E) together with the GH10 from *Paenibacillus* bin67. Other pioneering populations in the hemicellulose deconstruction process were *Thermobacillus* bin17 (C3), 51, 53 (C1 and C3), and *Paenibacillus* bin67 (Fig. 5E, cluster 1) through the expression of GHs from the families 10, 11, 16, 26, and 51 whose expression was significantly higher in SDM1 than in SDM3 *(p* < 0.01, log2fold > 1). The expression of these GHs was higher during Day 2 and then declined but continued through the incubation period. GHs that contributed to the high hemicellulose degrading activity in SDM1 were the GHs 10 and 16 from *Actinotalea* bin5 and *Rhosdospirillales* bin9, whose activity was detected since the beginning of the incubation and increased over time up to Day 14 (C1, Fig. 5E).

Significantly highly expressed GHs in SDM3 are shown in Fig. 3E cluster 2, and included GH10, 11, 16, 51, 53, and 67 from *Actinopolymorpha* bin90, *Mycobacterium* bin18, *Thermocrispum* bin46, and *Paenibacillus* bin42. The expression of these GHs increased over time with those from *Actinopolymorpha* bin90, *Mycobacterium* bin18, and *Thermocrispum* bin46 reaching higher levels from Day 9 to Day 14 likely indicating the critical roles of these populations for the progression of biomass decomposition in SDM3.

In contrast to pectin and hemicellulose, the expression of genes related to cellulose deconstruction was overall higher in SDM3 than in SDM1 (Fig. 5C). Expression patterns showed that the cellulose deconstruction commenced in both treatments (SDM1/SDM3) by the activity of *Actinotalea* bin1, *Gemmatimonadetes* bin10, *Paenibacillus* bin67, *Thermobacillus* bin53, and *Thermobacillus* bin51 expressing GH5 and GH9 (C1, Fig. 5F). In SDM3, the cellulose degradation process was complemented by the significantly higher activity *(p* < 0.01, log2fold > 1) of *Actinopolymorpha* bin90 expressing a GH5 and a GH9, *Thermocrispum* bin46 and *Salinispora* bin39 expressing an AA10, together with *Salinispora* bin64 expressing GH6, 9 and 48, all of which increased over time (C3, Fig. 5F). In SDM1, *Salinispora* bin32 was a key contributor to cellulose degradation through the expression of a GH5, 6 and 48 that reached its highest from Day 9 to Day 12. Other bacterial populations likely contributing to cellulosic activity were *Paenibacillus* bin34, *Dongia* bin26 (C5) and *Thermobacillus* bin17 (C2) through the expression of GH5 and GH9.

In comparison to bacterial polysaccharide deconstruction, bacterial lignin deconstruction is less understood^43^. Inspection of the metagenome and metatranscriptome identified a protein annotated as a multi-copper oxidase in the *Gemmatimonadetes* bin10. A homolog of this protein in a closely related thermophilic *Gemmatimonadetes* population was identified by proteomics as one of the most abundant proteins in the supernatant of bacterial consortium growing on switchgrass at 60 °C^44^. In addition, a homologous Cu-containing protein was identified in cultures of *Thermobifida fusca* growing on sugarcane bagasse^45^. This Cu-protein improved the polysaccharide hydrolysis of *T. fusca* glycoside hydrolases and improved the deconstruction efficiency of an engineered cellulosome on wheat straw when it was incorporated as a heterologous protein^46^. In the sorghum cultures, the *Gemmatimonadetes* bin10 multi-copper oxidase expression was found to be significantly higher in SDM3 than in SDM1 from Day 5 to Day 7, reaching similar levels at Day 9 (Figure 6). In addition, expression of a complete pathway for aromatic catabolism from 4-hydroxybenzoate transformation to protocatechuate and its conversion to succinyl-CoA and acetyl-CoA via the beta-ketopadipate pathway was observed in the *Filomicrobium* bin12. This pathway was detected at significantly higher levels in SDM3 compared to SDM1 from Day 2 to Day 7 (Fig. 6).

**Fig. 6.**
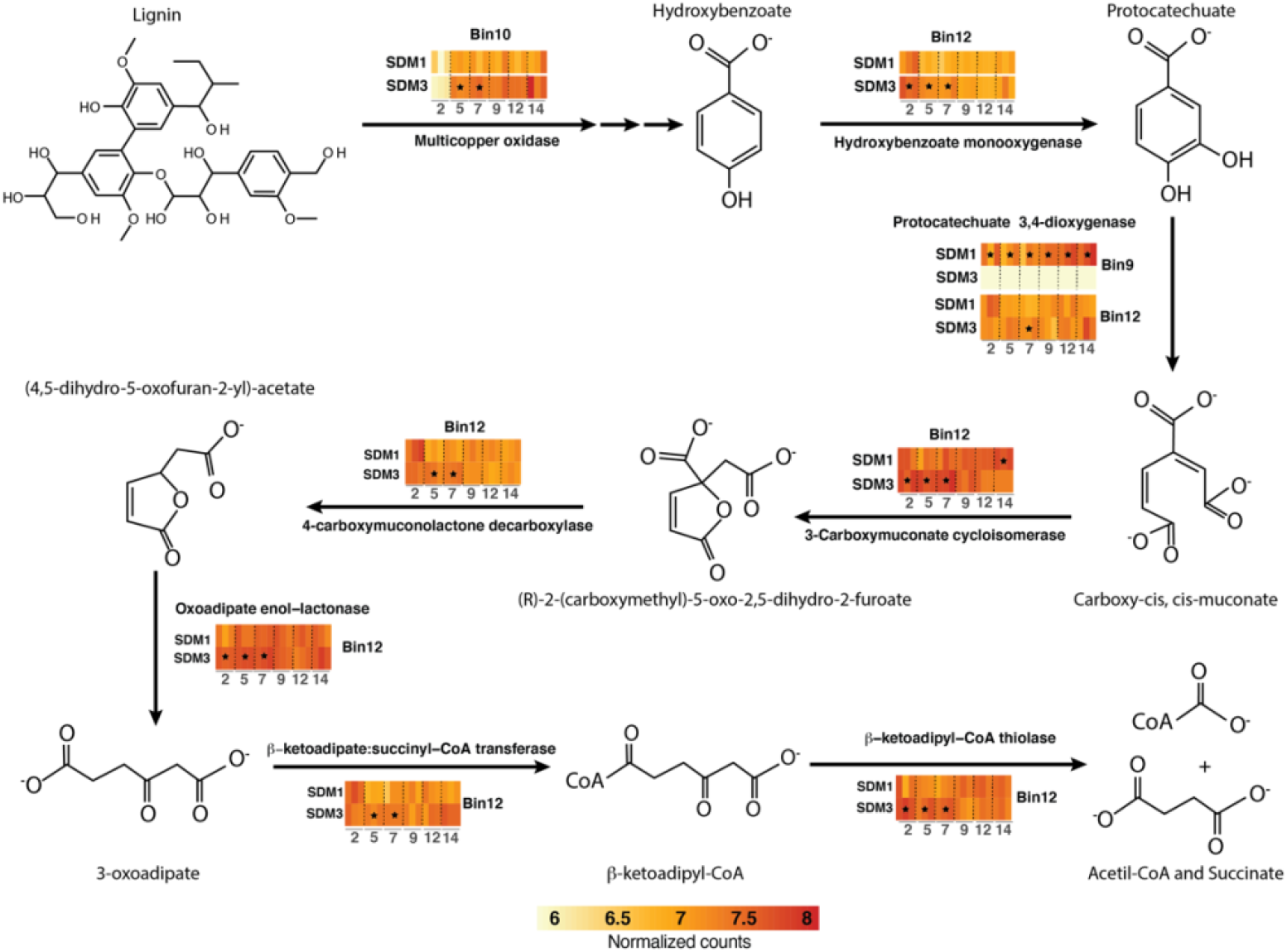
Schematic representation of the expression patterns for aromatic-degrading genes. The heatmaps are colored based on normalized counts for the targeted genes. Stars indicate the time points at which gene expression was significantly higher than in the opposite treatment *(p* < 0.01, log2fold > 1).

## Discussion

The two-tier cultivation of compost-derived microbiomes on sorghum led to the establishment of microbiomes for which community structure and performance could be assessed. Initial inoculation and growth on biomass sorghum provided distinct microbiomes *(comm 1-3)* that traversed independent trajectories during two months of adaptation (Fig. 1A). The development of distinctive microbiomes echoes parallel cultivation of microbiomes from *Sarracenia purpurea* pitcher plants grown on ground crickets^47^. The community structures of these parallel microbiomes also diverged during adaptation and the pitcher plant-derived consortia had variable activities in chitin deconstruction. The second-tier growth using the *comm 1-3* microbiomes as inoculum for growth on wild-type (SDM4-SDM6) and lignin-reduced sorghum varieties (SDM1-SDM3) demonstrated that the structure and deconstructive activities of these microbiomes are reproducible. This observation suggests that after adaptation the community structures are maintained, allowing detailed comparisons between microbiomes that are statistically robust. Furthermore, analysis of variance between our compost-enriched microbiomes grown on wildtype sorghum (SDM6) compared to the *bmr-6×12* mutant (SDM3) provides persuasive evidence that community structure, rather than plant cell wall structure, defines the trajectory of deconstruction. The increased digestibility of the *bmr-6×12* mutant is consistent with its reduced lignin content and resulting lower recalcitrance^24,41^.

Genome-resolved metagenomics demonstrated the most abundant populations in the microbiomes were two closely related *Actinotalea* populations. The most abundant *Actinotalea* population in the *comm1* and *comm3*-derived microbiomes *(Actinotalea* bin1) possessed fewer deconstructive enzymes than the most abundant *Actinotalea* population *(Actinotalea* bin5) in *comm2*; however, the performance of the *comm2*-derived microbiomes, as measured by biomass loss, cellulase/xylanase activity and lignin remaining in the residual biomass was generally lower compared to the *comm1* and *comm3-derived* microbiomes. The presence of *Gemmatimonadetes* bin10 and *Filomicrobium* bin12 in *comm1-3* and their daughter communities suggested their prominent role in biomass deconstruction. This hypothesis was confirmed by both network analysis of gene expression, which demonstrated that gene expression in these populations were correlated, and functional analysis, which demonstrated that the *Gemmatimonadetes* and *Filomicrobioum* populations were involved in lignin deconstruction, an essential function in the deconstruction of the secondary plant cell wall. In addition, *Paenibacillus* bin67 and *Thermobacillus* bin17, bin51 and bin53 are broadly distributed and demonstrated high, correlated expression of pectinases and hemicellulases, especially early in the two-week cultivation, that is consistent with deconstruction of the primary cell wall. The contribution of these lower abundance populations to cell wall deconstruction is a phenomenon which has been observed in native microbiomes that deconstruct complex polysaccharide substrates like the human gut^48^.

The microbiomes derived from *comm1* and *comm3* growing on the *bmr-6x12* sorghum mutant that were dominated by *Actinotalea* bin1 provided an opportunity to link the community performance, as measured by biomass loss and enzymatic activity, to detailed gene expression dynamics. Focusing on the genes for biomass deconstruction, the *comm1*-derived microbiome (SDM1), had higher levels of expression of pectin and hemicellulose deconstructing enzymes, with the peak of gene expression activity occurring during the initial time (Day 2) and the majority of genes being expressed by the *Firmicutes.* We interpret this pattern as initial deconstruction of the primary cell wall, which continues throughout the two-week cultivation. At Day 5, there was increased expression of the multi-copper oxidase from the *Gemmatimonadetes* bin10 population, consistent with the commencement of deconstruction of the secondary cell wall, and the relative expression level was higher in SDM3, the more active set of cultures. This increased expression was also mirrored in the aromatic catabolic genes expressed by *Filomicrobium* bin10, the majority of which were expressed from Day 2 to Day 7 at higher levels in SDM3. The cellulase genes, especially GH6, GH9, GH48 and AA10, are expressed by *Actinobacteria* (*Salinospora*, *Actinopolymopha, Thermocripsum*) later in the cultivation (Day 9 to Day 14) and at higher levels in SDM3. SDM1 and SDM3 form two separate clusters of cellulase expression, indicating that these activities are distinct between the two communities. This distinction is also seen in the network analysis, where *Salinospora* bin32 (SDM1) and *Salinopsora* bin64 (SDM3) are peripheral and divergent members of the network, suggesting the response to cellulose has less overlap between the two communities compared to the other plant polymers. The increases in gene expression are consistent with biochemical measurements which show SDM3 has higher cellulase activity. The observation of higher cellulase activity, which arises from the actinobacterial populations, may explain the increased biomass deconstruction by SDM3 communities. The overall pattern of community dynamics, with *Firmicutes* being active at early timepoints and Actinobacteria active and later timepoints, mirrors the dynamics observed during composting^49^.

The work described here highlights the importance of founder effects in defining the composition and trajectory of microbiomes, and reinforces the observation that subtle differences in community composition and the genomic content of strains may lead to significant differences in community performance^50^. These considerations should be accounted for in using microbiomes for biotechnology and building synthetic microbiomes^51^.

### Data availability

Metagenomic and metatranscriptomic sequencing data can be accessed at the Joint Genome Institute Genome Portal (http://genome.jgi.doe.gov/) under Proposal ID: 503813 (Alteration of lignin biosynthetic pathways in sorghum enhances its deconstruction by adapted microbial consortia).

## Acknowledgments

This work was performed as part of the DOE Joint BioEnergy Institute (http://www.jbei.org), supported by the US DOE, Office of Science, Office of Biological and Environmental Research, through contract DE-AC02-05CH11231 between Lawrence Berkeley National Laboratory and the US DOE. Metagenomic and metatranscriptomic sequencing was conducted by the Joint Genome Institute, which is supported by the Office of Science of the US DOE under contract no. DE-AC02-05CH11231. The forage sorghum biomass was provided by Scott Sattler at USDA Agricultural Research Station in Lincoln, Nebraska.

## Contributions

S. W.S and J.C.-N. designed experiments; L.M.T., M.A., K.D., Y.G., B.G.R. and A.E. performed experiments; L.M.T., M.A., Y.-W.W, N.X., J.C.-N. performed data analysis; C.L., J.C.M., P.A., T. R.N., H.S., B.A.S. supervised research; L.M.T., M.A., J.C-N. and S.W.S wrote manuscript. All authors approved the final manuscript.

## References

1. Cosgrove, D. J. & Jarvis, M. C. Comparative structure and biomechanics of plant primary and secondary cell walls. Front. Plant Sci. 3, 204 (2012).

2. Peralta-Yahya, P. P., Zhang, F., Del Cardayre, S. B. & Keasling, J. D. Microbial engineering for the production of advanced biofuels. Nature 488, 320–328 (2012).

3. Stephanopoulos, G. Challenges in engineering microbes for biofuels production. Science (80-.). 315, 801–804 (2007).

4. Kerr, R. A. Global warming is changing the world. Science (80-.). 316, 188–190 (2007).

5. Arfi, Y. et al. Characterization of salt-adapted secreted lignocellulolytic enzymes from the mangrove fungus Pestalotiopsis sp. Nat. Commun. 4, 1–9 (2013).

6. Straub, C. T. et al. Quantitative fermentation of unpretreated transgenic poplar by Caldicellulosiruptor bescii. Nat. Commun. 10, 1–6 (2019).

7. Blifernez-Klassen, O. et al. Cellulose degradation and assimilation by the unicellular phototrophic eukaryote Chlamydomonas reinhardtii. Nat. Commun. 3, 1–9 (2012).

8. Eibinger, M., Sattelkow, J., Ganner, T., Plank, H. & Nidetzky, B. Single-molecule study of oxidative enzymatic deconstruction of cellulose. Nat. Commun. 8, 1–7 (2017).

9. Himmel, M. E. et al. Biomass recalcitrance: Engineering plants and enzymes for biofuels production. Science (80-.). 315, 804–807 (2007).

10. Tan, T. C. et al. Structural basis for cellobiose dehydrogenase action during oxidative cellulose degradation. Nat. Commun. 6, 1–11 (2015).

11. Ceja-Navarro, J. A. et al. Gut anatomical properties and microbial functional assembly promote lignocellulose deconstruction and colony subsistence of a wood-feeding beetle. Nat. Microbiol. 4, (2019).

12. Gharechahi, J. et al. Metagenomic analysis reveals a dynamic microbiome with diversified adaptive functions to utilize high lignocellulosic forages in the cattle rumen. ISME J. 15, 1108–1120 (2021).

13. López-González, J. A. et al. Enzymatic characterization of microbial isolates from lignocellulose waste composting: Chronological evolution. J. Environ. Manage. 145, 137–146 (2014).

14. Romero Victorica, M. et al. Neotropical termite microbiomes as sources of novel plant cell wall degrading enzymes. Sci. Rep. 10, 3864 (2020).

15. Silva, P. C. et al. A novel d-xylose isomerase from the gut of the wood feeding beetle *Odontotaenius disjunctus* efficiently expressed in Saccharomyces cerevisiae. Sci. Rep. 11, 4766 (2021).

16. Jiménez, D. J. et al. Ecological Insights into the Dynamics of Plant Biomass-Degrading Microbial Consortia. Trends Microbiol. 25, 788–796 (2017).

17. Kolinko, S. et al. A bacterial pioneer produces cellulase complexes that persist through community succession. Nat. Microbiol. 3, 99–107 (2018).

18. Peng, X. et al. Genomic and functional analyses of fungal and bacterial consortia that enable lignocellulose breakdown in goat gut microbiomes. Nat. Microbiol. 6, 499–511 (2021).

19. Santos, J. et al. From nature to the laboratory: the impact of founder effects on adaptation. J. Evol. Biol. 25, 2607–2622 (2012).

20. Eudes, A. et al. SbCOMT (Bmr12) is involved in the biosynthesis of tricin-lignin in sorghum. PLoS One 12, (2017).

21. Eichorst, S. A. et al. Community dynamics of cellulose-adapted thermophilic bacterial consortia. Environ. Microbiol. 15, 2573–2587 (2013).

22. King, B. C., Donnelly, M. K., Bergstrom, G. C., Walker, L. P. & Gibson, D. M. An optimized microplate assay system for quantitative evaluation of plant cell wall-degrading enzyme activity of fungal culture extracts. Biotechnol. Bioeng. 102, 1033–1044 (2009).

23. Ing, N. et al. A multiplexed nanostructure-initiator mass spectrometry (NIMS) assay for simultaneously detecting glycosyl hydrolase and lignin modifying enzyme activities. Sci. Rep. 11, 1–9 (2021).

24. Sattler, S. E., Funnell-Harris, D. L. & Pedersen, J. F. Efficacy of Singular and Stacked brown midrib 6 and 12 in the Modification of Lignocellulose and Grain Chemistry. J. Agric. Food Chem. 58, 3611–3616 (2010).

25. Deng, K. et al. Rapid kinetic characterization of glycosyl hydrolases based on oxime derivatization and nanostructure-initiator mass spectrometry (NIMS). ACS Chem. Biol. 9, 1470–1479 (2014).

26. Barnes, W. & Anderson, C. Acetyl Bromide Soluble Lignin (ABSL) Assay for Total Lignin Quantification from Plant Biomass. BIO-PROTOCOL 7, (2017).

27. DeAngelis, K. M. et al. Strategies for enhancing the effectiveness of metagenomic-based enzyme discovery in lignocellulolytic microbial communities. Bioenergy Res. 3, 146–158 (2010).

28. Edgar, R. C. Search and clustering orders of magnitude faster than BLAST. Bioinformatics 26, 2460–2461 (2010).

29. Minh, B. Q. et al. IQ-TREE 2: New Models and Efficient Methods for Phylogenetic Inference in the Genomic Era. Mol. Biol. Evol. 37, 1530–1534 (2020).

30. McMurdie, P. J. & Holmes, S. Phyloseq: An R Package for Reproducible Interactive Analysis and Graphics of Microbiome Census Data. PLoS One 8, e61217 (2013).

31. Love, M. I., Huber, W. & Anders, S. Moderated estimation of fold change and dispersion for RNA-seq data with DESeq2. Genome Biol. 15, 550 (2014).

32. Prjibelski, A., Antipov, D., Meleshko, D., Lapidus, A. & Korobeynikov, A. Using SPAdes De Novo Assembler. Curr. Protoc. Bioinforma. 70, (2020).

33. Zhu, W., Lomsadze, A. & Borodovsky, M. Ab initio gene identification in metagenomic sequences. Nucleic Acids Res. 38, e132–e132 (2010).

34. Buchfink, B., Reuter, K. & Drost, H.-G. Sensitive protein alignments at tree-of-life scale using DIAMOND. Nat. Methods 18, 366–368 (2021).

35. Huson, D. H. et al. MEGAN Community Edition – Interactive Exploration and Analysis of Large-Scale Microbiome Sequencing Data. PLOS Comput. Biol. 12, e1004957 (2016).

36. Palumbo, A. et al. KBase: An Integrated Knowledgebase for Predictive Biology and Environmental Research. in Proceedings of the International Conference on Bioinformatics & Computational Biology (BIOCOMP) 1 (2014).

37. Aitchison, J. Principles of compositional data analysis. Lect. Notes-Monograph Ser. 73–81 (1994).

38. Zhou, J. et al. Functional Molecular Ecological Networks. MBio 1, (2010).

39. Zhou, J., Deng, Y., Luo, F., He, Z. & Yang, Y. Phylogenetic Molecular Ecological Network of Soil Microbial Communities in Response to Elevated CO2. MBio 2, (2011).

40. Shannon, P. et al. Cytoscape: a software environment for integrated models of biomolecular interaction networks. Genome Res. 13, 2498–504 (2003).

41. Godin, B. et al. Improved sugar yields from biomass sorghum feedstocks: Comparing low-lignin mutants and pretreatment chemistries. Biotechnol. Biofuels 9, (2016).

42. Mewis, K., Lenfant, N., Lombard, V. & Henrissat, B. Dividing the large glycoside hydrolase family 43 into subfamilies: A motivation for detailed enzyme characterization. Appl. Environ. Microbiol. 82, 1686–1692 (2016).

43. Brown, M. E. & Chang, M. C. Exploring bacterial lignin degradation. Curr. Opin. Chem. Biol. 19, 1–7 (2014).

44. D’haeseleer, P. et al. Proteogenomic Analysis of a Thermophilic Bacterial Consortium Adapted to Deconstruct Switchgrass. PLoS One 8, e68465 (2013).

45. Chen, C. Y., Hsieh, Z. S., Cheepudom, J., Yang, C. H. & Meng, M. A 24.7-kDa copper-containing oxidase, secreted by *Thermobifida fusca*, significantly increasing the xylanase/cellulase-catalyzed hydrolysis of sugarcane bagasse. Appl. Microbiol. Biotechnol. 97, 8977–8986 (2013).

46. Davidi, L. et al. Toward combined delignification and saccharification of wheat straw by a laccase-containing designer cellulosome. Proc. Natl. Acad. Sci. 113, 10854–10859 (2016).

47. Bittleston, L. S., Gralka, M., Leventhal, G. E., Mizrahi, I. & Cordero, O. X. Context-dependent dynamics lead to the assembly of functionally distinct microbial communities. Nat. Commun. 11, 1–10 (2020).

48. Flint, H. J., Scott, K. P., Duncan, S. H., Louis, P. & Forano, E. Microbial degradation of complex carbohydrates in the gut. Gut Microbes 3, 289–306 (2012).

49. Antunes, L. P. et al. Microbial community structure and dynamics in thermophilic composting viewed through metagenomics and metatranscriptomics. Sci. Rep. 6, 38915 (2016).

50. Thompson, A. W. et al. Robustness of a model microbial community emerges from population structure among single cells of a clonal population. Environ. Microbiol. 19, 3059–3069 (2017).

51. Strous, M. & Sharp, C. Designer microbiomes for environmental, energy and health biotechnology. Curr. Opin. Microbiol. 43, 117–123 (2018).

