## Supplemental Figures for "Assessing Comparative Microbiome Performance in Plant Cell Wall Deconstruction Using Multi-‘omics-Informed Network Analysis"

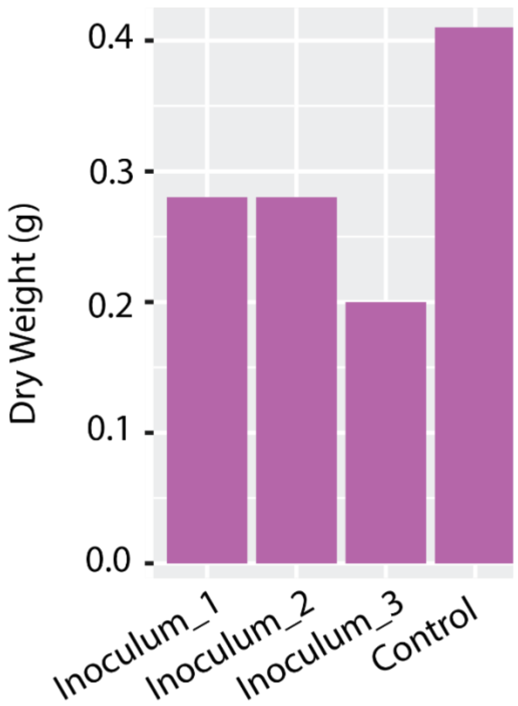


Supplementary Fig. 1. Residual dry weight biomass after a 56-day incubation. The figure shows that three incubations diverge in terms of biomass degradation even though the inoculum for the three samples was derived from the same 100-mg compost sample.

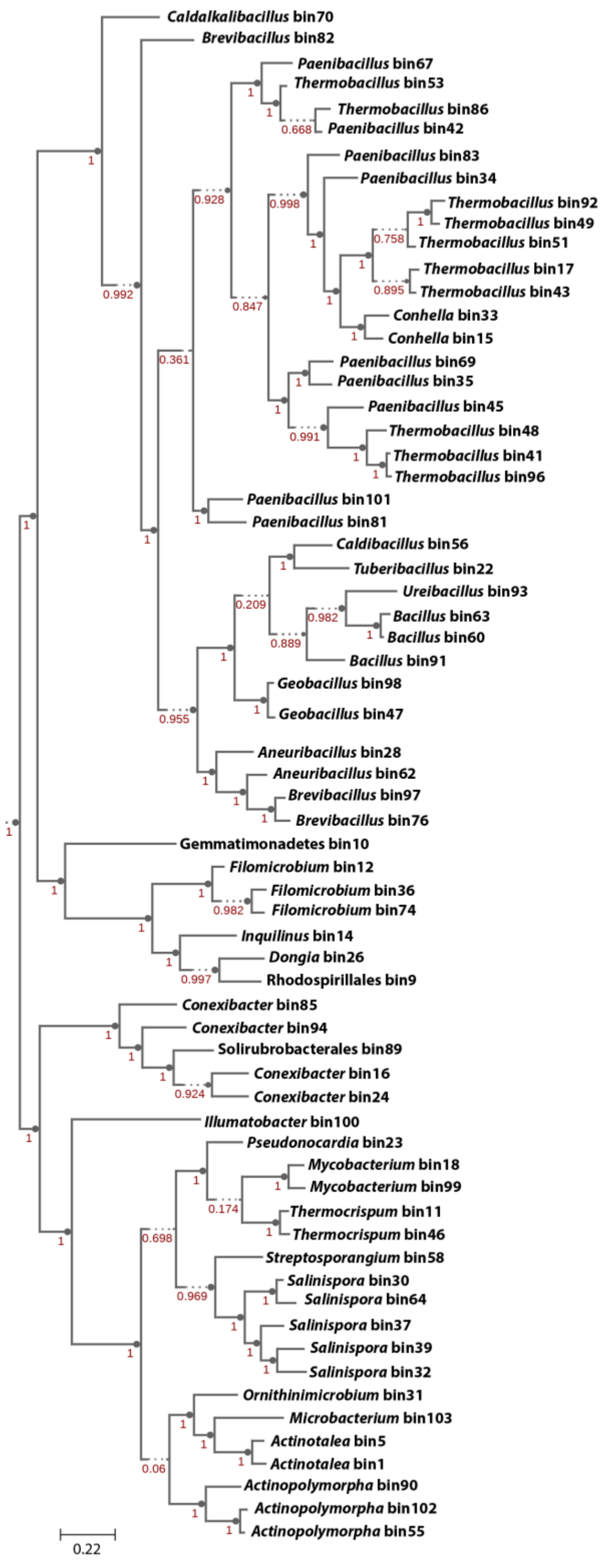


Supplementary Fig. 2. Phylogenetic tree reconstructed based on 49 core universal genes defined by Clusters of Orthologous Groups (COGs) using maximum likelihood. The tree was reconstructed on KBase.

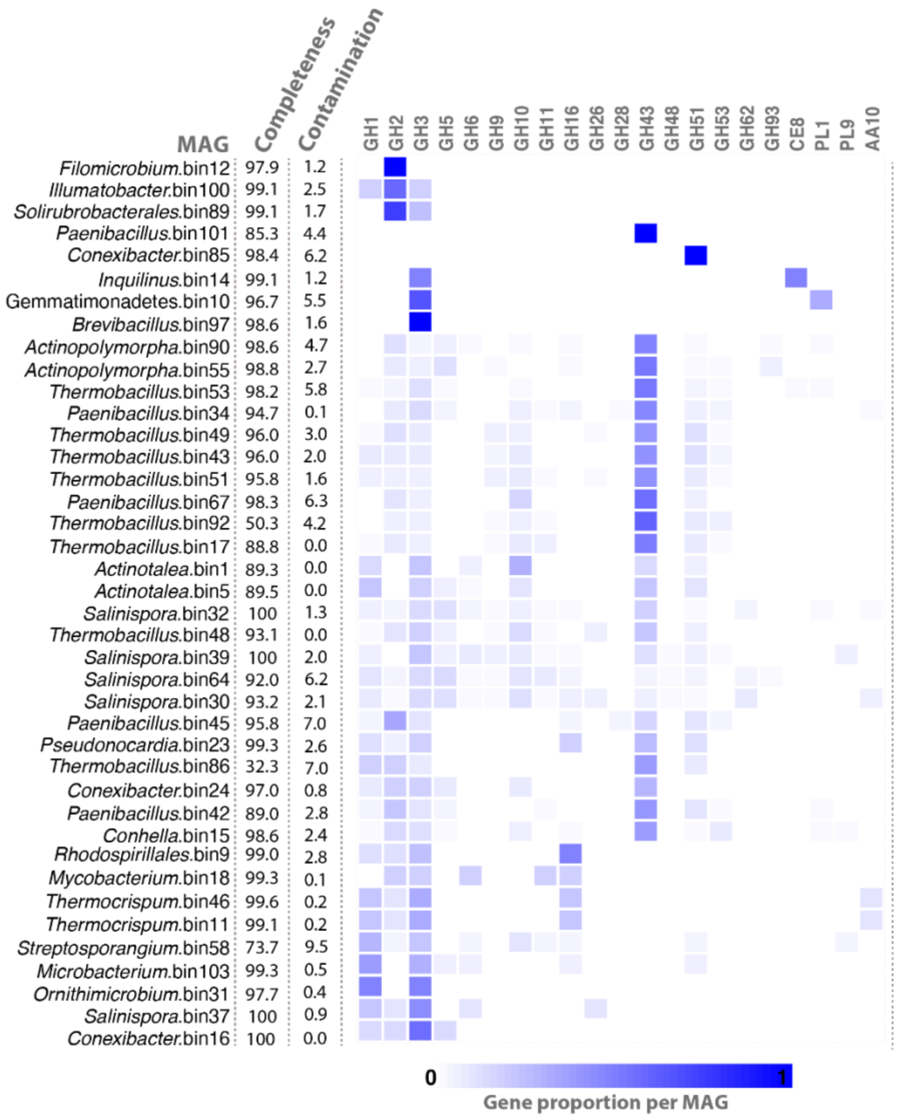


Supplementary Fig. 3. Gene proportion per MAG for selected GHs. MAGs with a completeness above 30% and contamination level lower than 10% are included in the figure.
